# Altered grid-like coding in early blind people

**DOI:** 10.1101/2023.05.19.541468

**Authors:** Federica Sigismondi, Yangwen Xu, Mattia Silvestri, Roberto Bottini

## Abstract

Spatial navigation in humans relies heavily on vision. However, the impact of early blindness on the brain navigation network and on the hippocampal-entorhinal system supporting cognitive maps, in particular, remains elusive. Here, we tested sighted and early blind individuals in both imagined navigation in fMRI and real-world navigation. During imagined navigation, the Human Navigation Network was reliably activated in both groups, showing resilience to visual deprivation. However, neural geometry analyses highlighted crucial differences between groups. A 60° rotational symmetry, characteristic of grid-like coding, emerged in the entorhinal cortex of sighted but not blind people, who instead showed a 4-fold (90°) symmetry. Moreover, higher parietal cortex activity during navigation in the blind was correlated with the magnitude of 4-fold symmetry and real-word navigation abilities. In sum, early blindness can alter the geometry of entorhinal cognitive maps, possibly as a consequence of higher reliance on parietal egocentric coding during navigation.

---

Humans heavily rely on vision to navigate the environment and acquire spatial information ^1–4^. However, the role of visual experience in the development and functioning of the brain system dedicated to spatial navigation remains largely unknown. In the brain, the human navigation network (HNN) comprises medial temporal, parietal, and occipital regions that support spatial memory and navigation across complementary allocentric and egocentric reference frames^5–11^. Significant breakthroughs in rodent electrophysiology attribute a crucial role in spatial navigation to the hippocampal-entorhinal system through the construction of allocentric cognitive maps^12^ based on the activity of highly specialized neurons, such as place cells^13^, head direction cells^14^ and grid cells^15^. Whereas place cells and head direction cells encode single locations and heading directions, respectively, grid cells in the medial entorhinal cortex (EC) present multiple firing fields and keep track of agent movements within the environment, tiling the navigable surface in a regular hexagonal grid. In addition to spatial navigation, hippocampal cognitive maps play a crucial role in representing the relationships between non-spatial stimuli. This general function in structuring information is central in human cognition^16, 17^ and supports the notion of a shared evolutionary link between the brain’s spatial navigation circuits and the human conceptual system^18, 19^. Compared to other modalities, vision provides highly precise and global information regarding the location of both proximal and distal landmarks, which are important for the creation of allocentric maps^20–22^, as well as compensating for cumulative error during path integration^23–25^. Notably, visual landmarks displaced in the environment as experimental manipulation cause a shift in the firing field orientation of grid, place, and head-direction cells toward the position of the cues, suggesting visual anchoring of cognitive maps^15, 26–29^. Consistently, when rodents need to navigate in the dark, after familiarizing themselves with an environment in plain light, both grid and head direction firing fields can be disrupted and navigation impaired^30, 31^. However, experiments with congenitally blind rodents have shown that place cell firing fields develop normally in the absence of vision, although with reduced firing rates^32^, and head direction cells maintain their directional selectivity but with lower precision^31^. Nothing is known about the resilience of place and head direction coding in blind humans, and the development of grid cells in the absence of functional vision remains untested across species. Notably, in the lack of vision, rodents can maintain stable allocentric maps of the environment by relying on olfactory cues^31^, a sense that, arguably, is not sufficiently developed in humans to serve the same purpose. How does early or congenital visual deprivation affect spatial navigation and its neural underpinnings in humans?

The ability to create mental maps of the environment is maintained in blind people^33, 34^, though early lack of vision can impair allocentric spatial coding^35^ (especially in large environments) and can lead to less reliable distance estimation^36^ and higher reliance on egocentric coding^34, 37^. A few neuroimaging studies have investigated the influence of early visual deprivation on the neural correlates underlying spatial navigation in early blind individuals (EB) using various tasks (e.g., sensory substitution, haptic and auditory navigation) and reporting inconsistent results^38–42^ (See discussion). Crucially, none of these experiments has investigated the impact of blindness on the hippocampal-entorhinal spatial codes. Given that EB seem to compensate for suboptimal allocentric representations by relying more on egocentric coding^22, 34, 43, 44^, we could hypothesize the existence in this population of a disruption of allocentric entorhinal spatial maps, together with selective recruitment of parietal spatial representations, which are reportedly more involved in egocentric processing of spatial information^45–47^. Indeed, one study has suggested that higher activity in the right inferior parietal cortex during spatial navigation correlates with ratings of navigational independence in EB^40^.

Here, in an fMRI experiment, we asked blindfolded sighted and EB participants to imagine navigating inside a clock-like space, walking from one number to another in a straight line, to investigate whether early lack of vision influenced the emergence of grid-like representations in the EB entorhinal cortex. The clock space was sampled at a granularity of 15° (24 unique paths) to provide a sufficient angle resolution to detect the hexadirectional signal (6-fold symmetry, Figure 1A), which is considered a reliable proxy for the activity of grid cells in fMRI both during visual^48–50^ and imagined navigation^51, 52^. Participants’ active navigation during the task was spurred by the introduction of a control question, solvable only by performing spatial inferences on the relationships between the positions of the different numbers inside the clock. Importantly, the clock environment was chosen because both EB and sighted participants are highly familiar with it and did not require extensive training; therefore, we benefited from this feature to control for a possible imbalance of task-familiarity between the two groups. We also ran a shorter, modified version of the experiment, in which clock navigation was compared to an arithmetic task (Nav-Math Experiment) to test possible differences between the sighted and blind in the activation of navigation-specific regions. Finally, we conducted a behavioral path-integration experiment to investigate the relationship between brain activations and spatial navigation abilities in both groups.

**Figure 1.**
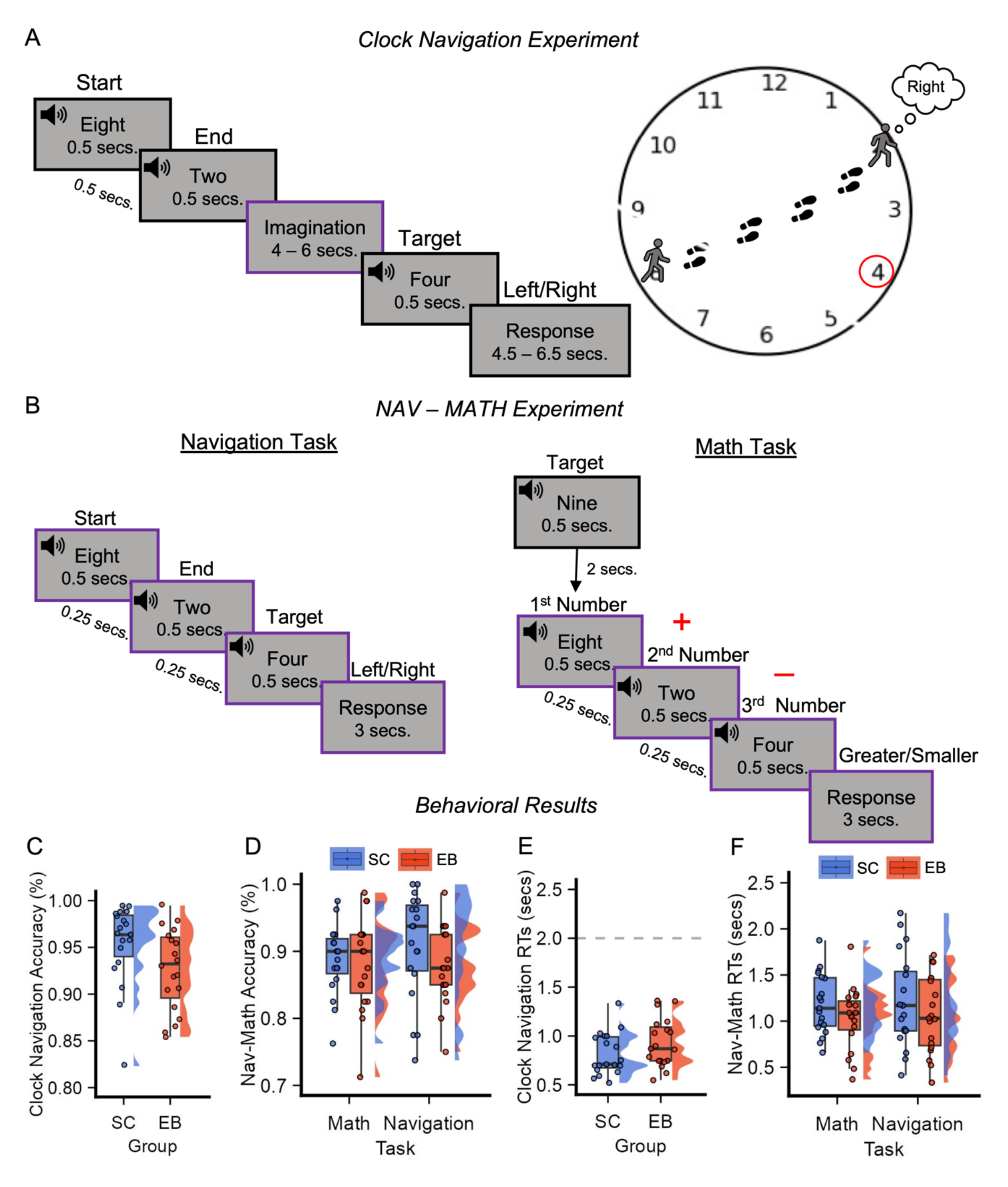
**Blind and sighted successfully navigated the clock space.** (A) Timeline of the Clock-Navigation experiment: participants were asked to imagine navigating from one number (Starting point) to another (Ending point) in the clock space, according to auditory instructions. Successful trials were denoted as those in which participants correctly indicated the position (left or right) of a third number (Target number) in the space, compared to their current trajectory, as shown in the graphical representation on the right. The purple box indicates the event of interest for the Quadrature Filter analysis. (B) Timeline of the Nav-Math experiment: the navigation task structure (left) was similar to that of the Clock Navigation experiment, with timing being the only difference. During the Math task (right), participants heard a number at the beginning of each math block (Target number) and then three more numbers. Participants were asked to sum the first two numbers, subtract the third number from that result, and then to compare the results with the target number. Purple boxes indicate the periods used for the univariate analysis (participant reaction times were used to determine the duration of the response period). (C) – (D) Accuracy of the two groups in the Clock-Navigation (C) and the Nav-Math (B) experiments: No difference in performance was detected between the two groups in the Clock-Navigation experiment (t(36) = 2.0, p = 0.052, two-tailed), nor between groups and tasks in the Nav-Math experiment (F(1,36) = 1.42, p = 0.24, two-tailed). (E) – (F) Reaction Times (RTs) of the two groups in the Clock-Navigation (E) and the Nav-Math (F) experiments: During the Clock-Navigation experiment, participants were required to indicate the position of the target number before a sound cue (with a response window of 2 secs). On average, all participants answered before the time limit (grey line), with no difference detected between groups (Chisq = 2.47, Df = 1, p = 0.11, two-tailed). In the Nav-Math experiment, participants were asked to give a response as quickly as possible. No differences were found between groups and tasks (Chisq = 0.24, Df = 1, p = 0.61, two-tailed).

## Results

### Blind and sighted successfully navigated the clock space

The imagined navigation task was similar across the two experiments (Clock-Navigation and Nav-Math), with the only exception in the timing (See STAR Methods and Figure 1A-B), and was developed by adapting imagined navigation tasks from previous studies^51, 52^. Blindfolded participants were auditorily provided with instructions concerning their starting and target locations (i.e., numbers) inside the clock and were instructed to think about themselves walking from one point to the other, taking the most straightforward path possible. After a jittered imagination period (4 – 6 secs), a third number was announced, and participants had to decide whether the number was located to their left or right. Successful trials were counted as those in which participants correctly responded to the control question. The arithmetic task (performed only in the Nav-Math experiment) was structurally similar to the navigation one. Participants heard three numbers, on which they had to perform simple operations (e.g., 8 + 2 - 4) and then compared the results to a target number presented at the beginning of each block (Figure 1B). Nineteen Early Blind (EB) and 19 matched Sighted Controls (SC) underwent an fMRI session, preceded by behavioral training in both tasks (math and navigation) and experiments (Nav-Math and Clock-Navigation).

Inside the MRI scanner, participants performed two runs of Nav-Math and eight runs of the Clock-Navigation experiments (see STAR Methods).

First, we computed the accuracy score for each participant and each experiment, examining the presence of differences between groups and/or tasks. No significant difference was detected between groups in the Clock-Navigation experiment (two-sample t-test; t_36_ = 2.0, p = 0.052, two-tailed, Figure 1C), nor between groups and tasks in the NAV-MATH experiment (Repeated-measure ANOVA; main effect of group: F_1,36_ = 0.86, p = 0.36; main effect of task: F_1,36_ = 0.85, p = 0.36; Group ×Task interaction: F_1,36_ = 1.42, p = 0.24, two-tailed; Figure 1D). Second, reaction times (RTs) of both groups were compared. In the Clock-Navigation experiment, no differences in RTs were detected between groups (Linear Mixed-Effect Model; Chisq = 2.45, Df = 1, p = 0.12, two-tailed, Figure 1E). Similarly, there were no differences in RTs between groups and tasks in the NAV-MATH experiment (Linear Mixed-Effect Model; main effect of group: Chisq = 1.53, Df = 1, p = 0.21; main effect of task: Chisq = 3.59, Df = 1, p = 0.06; Group × Task interaction: Chisq = 0.24, Df = 1, p = 0.62, two-tailed, Figure 1F). Collectively, these results suggest that both sighted and blind were able to perform the tasks and to perform comparably.

### The human navigation network is resilient to early visual deprivation

The brain network typically activated during spatial navigation tasks (Human Navigation Network; HNN) comprises frontoparietal, occipital, and medial temporal lobe regions^53–60^. Here, we investigated whether early blind and sighted populations rely on similar neural networks during imagined navigation in the clock space.

We analyzed fMRI data from the Nav-Math experiment to detect the brain regions with a greater level of activation during navigation compared to mathematics using a brain mask comprising brain regions in the Human Navigation Network (HNN) downloaded from Neurosynth (Figure S2, see STAR methods). The mask included (bilaterally) the middle frontal gyrus (MFG), superior parietal lobe/precuneus (SPL), retrosplenial cortex (RSC), occipital place area (OPA), parahippocampal place area (PPA) and hippocampus. Importantly, the navigation and mathematic task used the exact same numerical stimuli (see Figure 1B) with different instructions.

The univariate contrast (i.e., Navigation > Math) revealed significant activity in all the ROIs, except for the hippocampus, in both sighted and blind (Figure 2B-C and Table S3, all results thresholded at p_FDR_ < 0.05). No significant cluster of activation emerged when we contrasted the two groups ([Navigation > Math] × [EB vs. SC]). Highly similar results emerged from unmasked whole-brain analysis (Figure S3). Given that previous studies have found significant differences between SC and EB during navigation tasks using univariate contrasts^38, 39^, we performed a whole-brain analysis to investigate whether some differences emerged across groups in regions that were not included in our HNN mask. This analysis did not reveal any significant difference between groups when correcting for multiple comparisons (p_FDR_< 0.05). This null result could be due to a lack of power (the Nav-Math Experiment consisted of just two runs), high variability across subjects, or both. However, previous studies report that blind individuals tend to rely more on egocentric coding of space compared to sighted individuals^34, 35, 37^. Therefore, to investigate if EB participants recruit parietal-based egocentric representation during navigation more than SC, we conducted a Small Volume Correction (SVC) analysis within the inferior parietal cortex (IPC), a region that has been implicated in the egocentric coding of spatial and non-spatial information^19, 61^ and spatial neglect^62, 63^. We used a spherical ROI (Radius = 10mm) centered on Montreal Neurological Institute (MNI) peak coordinates (36/-68/44) obtained from an independent study investigating egocentric spatial representations during imagined navigation in a circular environment, structurally similar to our clock^47^. The analysis revealed a significant difference during imagined navigation between SC and EB within the inferior parietal cortex ([Navigation > Math] × [EB > SC]; SVC: Voxel level p < 0.001, Cluster-level p_FWE_ < 0.05; see STAR METHODS and Figure S3). Supporting the SVC analysis, whole-brain results using a lenient threshold (p < 0.005 uncorrected) revealed the emergence of a cluster of activation in the right inferior parietal cortex (IPC, 39/-61/53, Figure 2D), overlapping with our ROI. We then used this inferior parietal cluster as an independently localized ROI for analysis in the Clock-Navigation experiment to test for possible parietal-based compensation during navigation in the blind group.

**Figure 2.**
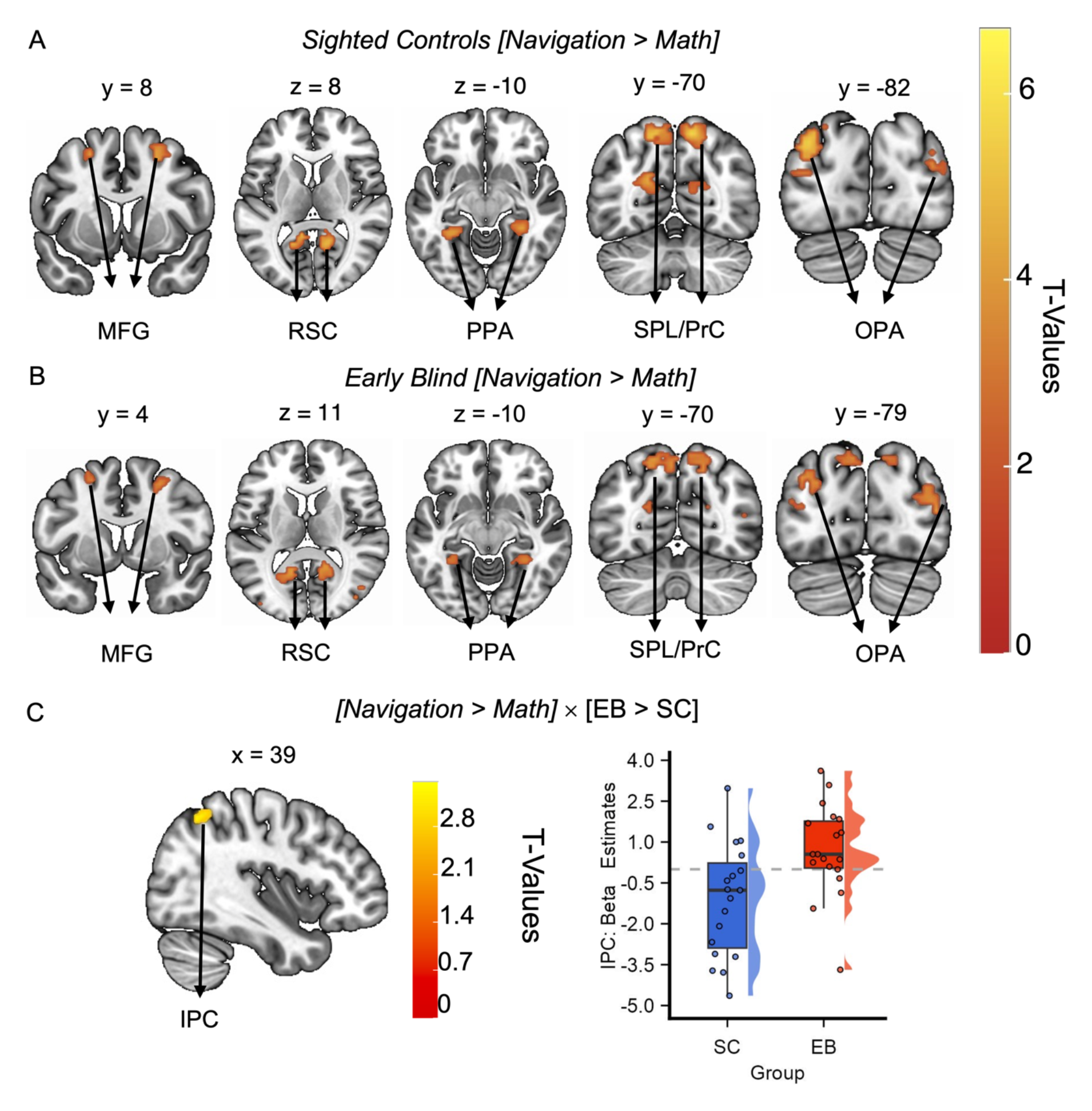
**The human navigation network is resilient to early visual deprivation.** (A) – (B) Results of the univariate analysis [Navigation > Math] performed within each group on the Nav-Math experiment data: the results showed activation in both SC (B) and EB (C) participants in the HNN mask (the activations were thresholded at p_FDR_ < 0.05 and overlapped with the MNI-152 T1 template) (C) Whole-brain (left), group-level univariate analysis ([Navigation > Math] × [EB > SC]) on unmasked data revealed higher activation in the inferior parietal cortex in EB compared to SC individuals when observed using a lenient threshold (IPC, 39/-61/53, p < 0.005 uncorrected, activation overlapped with the MNI-152 T1 template). Right: beta estimates extracted within the IPC cluster (Navigation > Math) for each group confirmed that the parietal cortex was more involved in the navigation task in the EB group.

In sum, during imagined navigation, EB and SC activated largely the same regions of the human navigation network without significant differences. To our knowledge, this is the first time that the HNN has been selectively detected in early blind during a navigation task, indicating that such a network is largely resilient to early visual deprivation.

### Six-fold grid-like coding did not emerge in early blind

Next, we analyzed the fMRI data from the Clock-Navigation Experiment to investigate whether a grid-like representation could be detected in the entorhinal cortex (EC) of both the sighted and early blind. In fMRI, grid-like coding can be detected as a 60° sinusoidal modulation of the Blood-Oxygen-Level-Dependent (BOLD) signal (6-fold symmetry), elicited by spatial trajectories aligned with the main axis of the hexagonal grid^48^. For this purpose, we applied a four-way cross-validation procedure and implemented quadrature filter analysis in which three partitions (six runs) of the data were used to estimate the subject-specific grid orientation (phase, ϕ), and the remaining partition (two runs) was used to test the strength of the 60° sinusoidal modulation of the BOLD signal in EC after realigning the trajectories’ angles to each participant’s specific grid orientation^48, 49^ (see STAR methods). Considering the lack of an a priori hypothesis on grid-coding lateralization, we first combined all sighted control bilateral ECs into a single ROI, thus estimating a common grid orientation for the left and the right EC^49^. No significant 60° sinusoidal modulation (6-fold symmetry) of the BOLD signal was found in the sighted bilateral EC (One sample t-test; t_18_ = 1.07, p = 0.30, two-tailed). We then performed a test for hexadirectional coding in the two hemispheres separately, and we found a significant signature of grid-like coding in the sighted control left EC (Bonferroni-corrected for multiple comparisons across hemispheres, one-sample t-test; t_18_ = 2.71, p = 0.014, one-tailed, Figure 3C) but not in the right EC (Bonferroni-corrected for multiple comparisons across hemispheres; one-sample t-test; t_18_= - 0.04, p = 0.5, one-tailed; Figure 3C). Equal analyses performed on alternative models in the left EC (4-,5-,7-, and 8-fold symmetry) confirmed the specificity of our 6-fold symmetry results (Bonferroni-corrected across hemispheres; one-sample t-test; 4-fold: t_18_ = - 1.58, p = 0.26; 5-fold: t_18_ = 0.35, p = 0.73; 7-fold: t_18_ = 1.38, p = 0.18 and 8-fold: t_18_ = −0.73, p = 0.47, all results one-tailed, Figure 3C). The same analysis was conducted on the blind population. Accounting for the non-normal distribution of 6-fold symmetry beta estimates in this group (Shapiro-Wilk normality test; Bilateral EC: W = 0.87, p = 0.01; Left EC: W = 0.89, p = 0.03 and Right EC: W = 0.81, p = 0.001, see STAR methods), we performed a Wilcoxon signed-ranked test instead of Student’s t-tests. We did not find a significant 6-fold symmetry in the EB group, neither when considering the bilateral EC ROI (V= 77, z = −0.7 p = 0.24, one-tailed) nor the hemispheres separately (Left EC: V= 86, z = −0.34; p = 0.37 and Right EC: V= 104, z = - 0.34; p = 0.37, all results one-tailed; Figure 3D). Notwithstanding the inconsistent pattern of results found in the two groups, grid-like representation in the early blind left EC was not significantly lower than the one expressed by sighted controls (Wilcoxon ranked-sum test W = 231, z = −1.46, p = 0.14, two-tailed, Figure 3E). This non-significant result might be attributed to the higher individual differences observed within the EB group (Standard deviation from the mean (SD); SC = 0.21; EB = 0.96; Levene’s test for Homogeneity of variance between groups; F_1,36_ = 11.69, p = 0.001; two-tailed).

**Figure 3.**
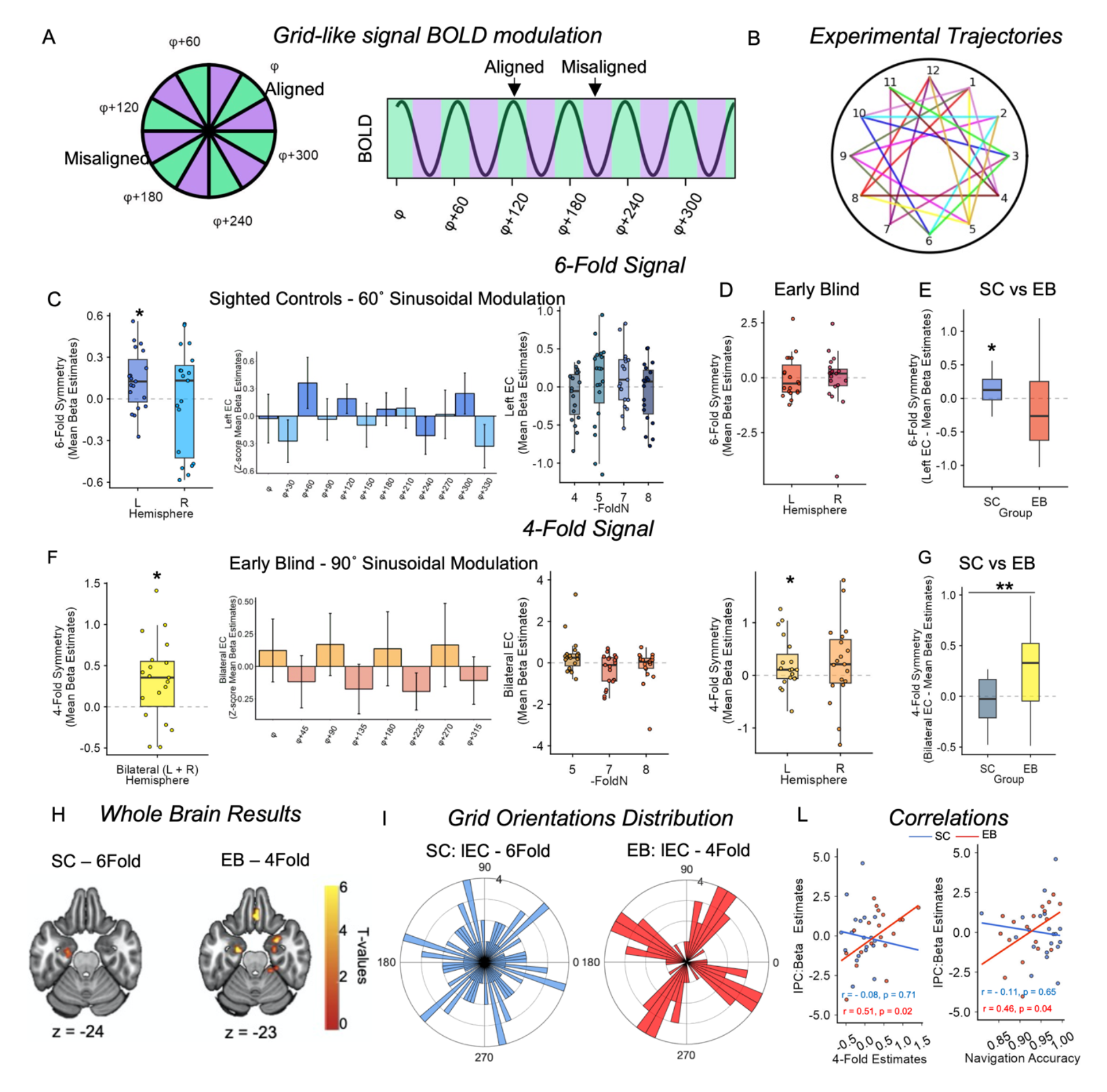
**Early blind and sighted showed different neural geometries in the entorhinal cortex.** (A) Left: grid cells fire more when the navigator moves in a direction aligned (green) with the preferred grid orientation (ϕ) and its 60° multiples, compared to misaligned directions (purple). Right: grid-like coding can be detected using fMRI as a 60° sinusoidal modulation of the BOLD signal (6-fold symmetry), as the conjunctive activation of grid cells produced a higher BOLD modulation for movements aligned rather than misaligned with the main grid orientation. (B) Participants performed 24 trajectories in each run, sampling the space with a granularity of 15°. (C) – (D) Quadrature filter analysis revealed the presence of a significant 6-fold symmetry in the sighted control left EC (C, left and middle panel, Bonferroni-corrected, one sample t-test; t_18_ = 2.71, p = 0.014 one-tailed). No significant modulations of the BOLD signal by the alternative-tested periodicities were detected (right panel). No 6-fold symmetry modulation was detected in the EB EC (D, Wilcoxon signed-rank test; Left EC: V= 86, z = −0.34; p = 0.37 and Right EC: V= 104, z = −0.34; p = 0.37, all results one-tailed). (E) No differences between groups were observed (Wilcoxon ranked-sum test W = 231, z = −1.46, p = 0.14, two-tailed), despite the detection of a reduced grid-like signal in the EB left EC. However, the highly variable distribution of 6-fold symmetry beta estimates in EB might have influenced the result. (F) Quadrature filter analysis performed on alternative periodicities (90°: 4-fold; 72°: 5-fold; 51.4°: 7-fold and 45°: 8-fold) revealed a significant 90° sinusoidal modulation of the BOLD signal (F, right and middle panel) in EB Bilateral EC (Bonferroni-corrected across four periodicities, one-sample t-test; p = 0.025, one-tailed). No other significant modulations of the BOLD signal were detected (right panel). (G) The 4-fold symmetry effect was significantly higher in early blind compared to sighted control participants (t_36_ = −3.18, p = 0.003, two-tailed). (H) Whole-brain, group-level analyses revealed a cluster of activation in the left EC of sighted and bilateral EC of early blind participants for the periodicity of interest (60° and 90° respectively; The activation overlaps with the MNI-152 T1 template at an uncorrected threshold of p < 0.01), for display only. (I) Sighted control 6-fold symmetry, left EC grid orientations were uniformly distributed in space (left, Bonferroni-corrected, Rayleigh’s test for circularity uniformity; p = 1), in line with previous findings in non-polarized environments. Surprisingly, the 4-fold symmetry grid orientations in the left EC of early blind individuals were significantly clustered (right, Bonferroni-corrected, Rayleigh’s test for circularity uniformity; p = 0.018). (L) Inferior Parietal Cortex activity during navigation in the Clock navigation experiment (Navigation > rest, separately for each group) significantly correlates both with the magnitude of the 4-fold symmetry effect (r = 0.51, p = 0.02, left panel) and with the accuracy in the Clock Navigation experiment (r = 0.46, p = 0.04, right panel) in EB. No correlations were found in SC participants. The beta estimates were extracted using the IPC cluster (39/-61/53) found during previous analysis as an independent ROI ([Navigation > Math] × [EB > SC] in the NAV-MATH experiment).

### A different neural geometry in the blind’s entorhinal cortex

Given the absence of 6-fold symmetry in the EC, we tested whether EB showed a significant difference in one of the alternative control models. This analysis revealed the emergence of a significant 90° sinusoidal modulation of the BOLD signal (4-fold) in the EB bilateral EC (Bonferroni-corrected across four tested periodicities, one-tailed, one-sample t-test; t_18_ = 2.77, p = 0.025, Figure 3F) and no other significant sinusoidal modulations (Bonferroni-corrected across four tested periodicities, one-tailed, one-sample t-test; 5-fold: t_18_ = 1.72, p = 0.2; 7-fold: t_18_ = - 1.95, p = 0.13, and 8-fold: t_18_ = - 0.89, p = 0.76, Figure 3F). Crucially, the magnitude of 4-fold symmetry in EB was significantly higher than SC (Two-sample t-test; t_36_ = −3.18, p = 0.003, two-tailed; Figure 3G). The 4-fold effect in EB was also present, although weakened, when we considered each hemisphere separately (One-sample t-test; Left EC: t_18_= 1.88, p = 0.038; Right EC: t_18_= 1.36, p = 0.095, one-tailed). Then, we further investigated the emergence of the 4-fold symmetry in EB, based on related findings in the literature. Entorhinal 4-fold symmetry in humans has already been observed in two studies, in which sighted participants navigated a virtual reality environment^64, 65^. Notwithstanding the differences in the environmental settings of the two studies (compartmentalized environment vs. barrier-free environment), they both attribute the emergence of the 90° sinusoidal BOLD modulation in the EC to the influence exerted by cardinal axes (north-south, east-west) in guiding the navigation behavior.

Interestingly, the 4-fold symmetry has been reported to be accompanied by a clustering of the preferred grid orientation (estimated for each participant) along the major axes of the environment^64^, suggesting the anchoring of the entorhinal map to these perpendicular axes. Thus, we tested whether a similar clustering was taking place for the blind in our experiment, considering that in a circular environment, no clustering is expected since there are no regular boundaries for anchoring^48, 49^. First, we looked at the phase distribution in the left EC in SC for 6-fold symmetry, and we indeed found that they were uniformly distributed (Bonferroni-corrected across hemispheres, Rayleigh test of Uniformity; p = 1, two-tailed, Figure 3I). However, the phases of the 4-fold symmetry in EB in the same hemisphere were significantly clustered (Bonferroni-corrected across hemispheres, Rayleigh test of Uniformity; p = 0.018, two-tailed, Figure 3I). Although this effect was not present in the right hemisphere, the highly significant clustering effect found in the left EC, consistent with the previous report of 4-fold deformation in humans^64^, suggests that the emergence of this neural geometry can be related to anchoring behavior on the main axes of the environment.

In a recent study, the disruption of 6-fold symmetry in the EC has been related to the adoption of an egocentric perspective during navigation and the increased activity of inferior parietal areas^66^. Since blind individuals tend to adopt mostly egocentric perspectives during navigation, might the emergence of the 4- fold symmetry be related? We explored this possibility by extracting the beta estimates obtained during a univariate whole-brain analysis, contrasting the navigation period to the resting period, independently for each group^67^ (see STAR method), using the inferior parietal cortex cluster obtained during the analysis of the Nav-Math experiment (defined in the contrast ([Navigation > Math] × [EB > SC], see above) as an independent ROI. A significant positive correlation emerged between 4-fold symmetry estimates and the activity in the IPC in EB (Pearson’s product-moment correlation; r = 0.51, p = 0.024, Figure 3L) that survived when controlling for 6-fold symmetry estimates (partial correlation: r = 0.49, p = 0.037). The same effect did not emerge in SC (Pearson’s product-moment correlation; r = −0.089, p = 0.71, Figure 3F). Interestingly, the activity of IPC in EB also correlated with their accuracy during the Clock-Navigation experiment (r = 0.46, p = 0.48; Partial Correlation 4-fold: r = 0.53, p = 0.02, Fig. 3I), whereas this correlation did not emerge in SC (r = −0.11, p = 0.65, Fig. 3L).

These results show that 6-fold symmetry in the EB entorhinal cortex was reduced, or at least more variable, than in SC during imagined navigation. Importantly, a different entorhinal geometry emerged in EB, one characterized by a 4-fold modulation absent in SC. The 4-fold entorhinal grid in EB was anchored to the main axes of the clock environment (in line with the emergence of 4-fold symmetry during spatial navigation in previous literature). Moreover, the strength of 4-fold symmetry in EC correlated with the strength of inferior parietal activity during navigation in EB, consistent with previous findings reporting a possible influence of the adoption of egocentric strategies on entorhinal grid-like coding^65, 66^.

### Compared to the sighted, early blind rely more on egocentric navigation strategies during everyday activities and performed worse in a path integration task

In order to assess their everyday navigation strategies, as well as their navigation abilities, we administered to sighted and blind participants a self-report questionnaire for navigation strategies^68^ and a path integration (PI) task. The questionnaire scores were first used to compute a general “Navigation Confidence” score for each participant. Second, we isolated the questions related to the use of Survey and Route knowledge, associated with Allocentric and Egocentric space representations, respectively^69–74^, and computed an independent score for each. The preference to rely more on one of the two strategies was assessed by computing the difference between Route and Survey scores (dRS); a positive value indicates participant preference to rely on Route strategies, whereas a negative value indicates the opposite (see STAR methods). No difference between the two groups was detected in the expressed confidence in their spatial navigation abilities (Two-sample t-test; Navigation Confidence score: t_36_ = −0.77, p = 0.44, two-tailed; Figure 4A). However, when considering the dRS, a significant difference between the two groups emerged (Wilcoxon ranked-sum test: W = 94, z = −2.52, p = 0.01, two-tailed; Figure 4B), indicating EB preference for Egocentric strategies during everyday navigation. This was also confirmed by a qualitative assessment of EB and SC strategies for navigating in the clock space (see STAR Methods), which indicates a preference of EB to adopt more egocentric-related strategies (e.g., imagining rotating the clock so that the starting and ending point were aligned with the body midline), compared to SC, which reported more allocentric-related strategies (See Table S4).

**Figure 4.**
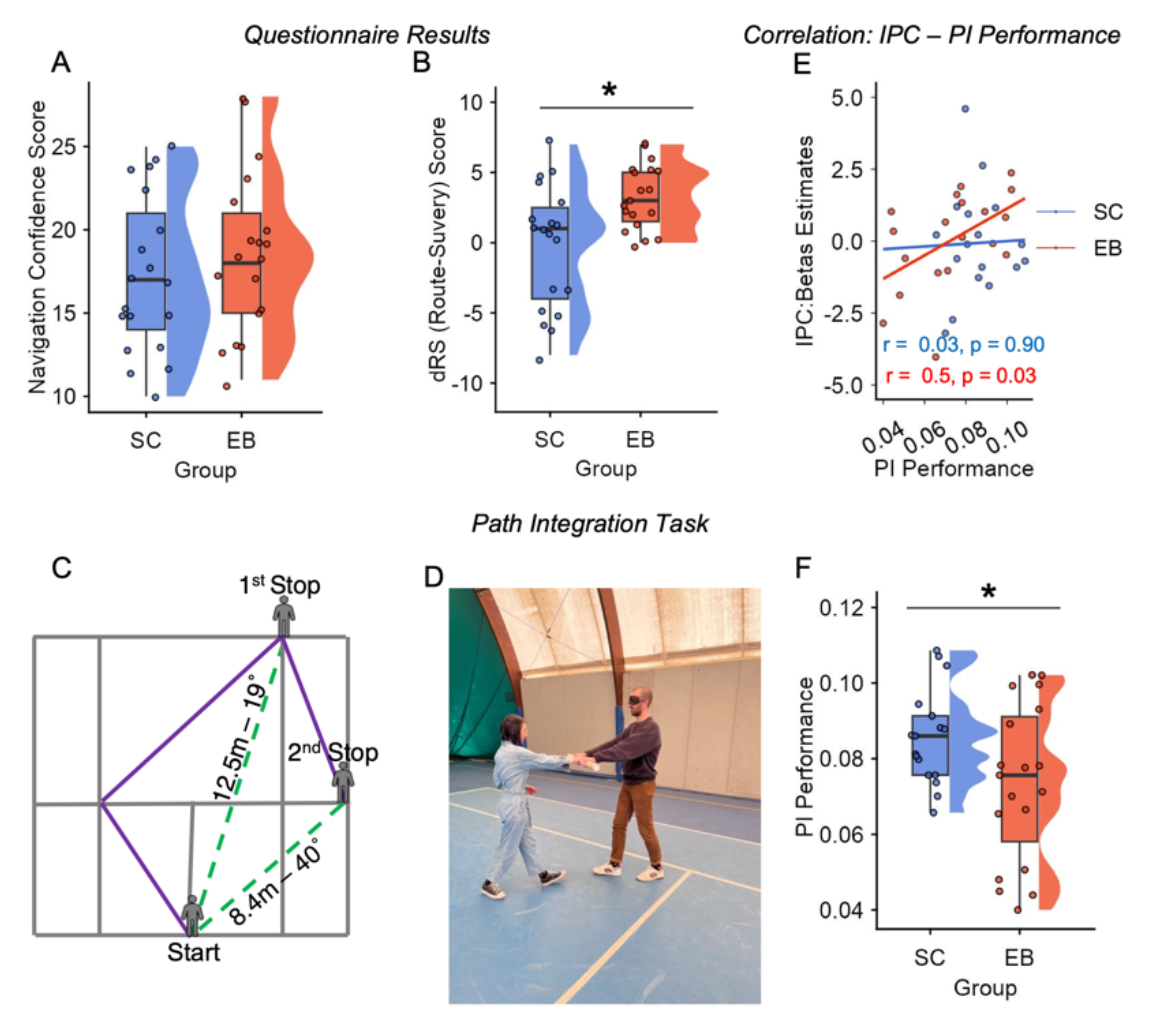
**Compared to SC, EB relied more on egocentric navigation strategies during everyday life and performed worse in a path integration task.** (A) – (B) Participants were asked to fill out a self-report questionnaire on their navigation abilities during everyday life. No differences between groups were detected concerning confidence in spatial navigation abilities (A). However, when differentiating between Route and Survey knowledge, we observed a significant difference between the two groups, with EB reporting a greater reliance on Route strategies than SC (B, W= 94, p = 0.01). (C) Path integration task: Half of a tennis court (10m × 11m) was used to perform the experiment. The experimenter led the blindfolded participants through several paths (purple lines), created using tennis court lines as a reference. Each path contained two stopping points, at which participants were required to estimate the distance and orientation between their actual position and the starting position (green dashed lines). (D) During the path integration experiment blindfolded participants were led by the experimenter trough the different paths as shown in the picture. (E) Inferior parietal cortex activity positively correlated with PI Performance in EB (r = 0.5, p = 0.03) but not SC (r = 0.03, p = 0.90). (F) PI performance for each participant was computed as 1/PI Error. PI Errors were calculated, integrating orientation and distance estimations as the Euclidean distance between the estimated and the real starting point positions. Results showed that SC were overall more accurate, compared to EB (t34 = 2.10, p = 0.04, two-tailed).

Path integration, defined as the ability to rely on idiothetic cues to update one’s current position in the environment in relation to the previously computed position^75, 76^, has been reported to be a sensitive hippocampal-entorhinal based test for allocentric spatial memories^77, 78^. We thus performed a Path Integration experiment on the same participants who performed the MRI task to investigate whether performance in a real-world navigation task was worse in EB.

Blindfolded participants (17 SC and 19 EB) walked along 16 linear paths with the help of an experimenter (see Figure S1 and Table S2). Each path contained two stopping points, at which participants were asked to estimate the distance and the orientation between their actual position and the starting point (Figure 4A). PI performance was calculated for each participant by combining their accuracy in estimating distances and orientations^65^ (see STAR methods). Our results showed that, relative to SC, EB PI performance was significantly lower (Two-sample t-test; t_34_ = 2.10, p = 0.04, two-tailed, Figure 4C).

### Parietal cortex activity predicted real-world navigation performance in early blind

Right inferior parietal cortex activity has recently been associated with the accuracy with which people encode spatial information, especially when environmental settings or experimental tasks enhance the use of egocentric, rather than allocentric, spatial representations^65, 66^. In line with this evidence, in the Clock-Navigation experiment, we observed a positive correlation in EB (who tend to use more egocentric related strategies to navigate) between right inferior parietal cortex (IPC) activity and accuracy during navigation (see above). Would IPC relative activation also correlate with navigation accuracy in the real world? To answer this, we correlated the beta estimates extracted in the right IPC during navigation in the Clock-Navigation experiment with accuracy in the PI experiment. We found a positive correlation between IPC activity and accuracy during the PI task in EB (r = 0.50, p = 0.028, Figure 4E), which was largely preserved when we controlled for 6-fold symmetry estimates (partial correlation: r = 0.48, p = 0.041) and 4-fold symmetry estimates (partial correlation: r = 0.45, p = 0.05). By contrast, no significant correlation was observed in SC between IPC activity and real-word navigation (r = 0.031, p = 0.91; Figure 4E), nor between 4-fold or 6-fold symmetry and PI performance (4-fold symmetry: r = 0.049, p = 0.85; 6-fold symmetry: r = −0.19, p = 0.44). Similarly, we did not observe a significant correlation between PI performance and 4-fold symmetry in EB (r = 0.24, p = 0.33), although we detected a negative correlation between 6-fold symmetry estimates and PI performance that was, however, driven by outliers and therefore should be taken with caution (6-fold symmetry: r = −0.5, p = 0.03, see supplementary materials and Figure S5). This result suggested that, in EB, the activity of parietal cortex is associated with better path integration abilities.

## Discussion

In the current study, we investigated the influence of early visual deprivation on the development of the HNN as well as the stability of grid-like coding in the hippocampal-entorhinal system. Here, we discuss the two main findings.

First, the HNN was reliably and selectively activated in both SC and EB during imagined navigation. Results of the Nav-Math experiment indicate that both sighted and blind participants have greater activation (bilaterally) in the medial frontal gyrus (MFG), retrosplenial cortex (RSC), parahippocampal place area (PPA), superior parietal lobule/precuneus (SPL/Precu) and occipital cortex (OPA) in the navigation task compared to the arithmetic task. We did not find navigation-specific activity in the hippocampus, in either SC or EB. This can be explained by the familiarity of both groups with the clock environment, which would not have demanded a strong reliance on episodic memory to retrieve its spatial configuration^79^.

Two previous studies have tested SC and EB performing spatial navigation during fMRI using a sensory substitution device based on tongue stimulation^38^ or the haptic navigation of a 3D maze^39^. In both cases, early blind people activated part of the HNN consisting of PPA, MFG, and SPL/Precu. Activity in the parahippocampal regions was stronger in EB than in SC, and in general, EB had higher occipital activations (lingual gyrus, cuneus, fusiform gyrus) than matched sighted controls. By contrast, we did not find any difference between groups in the HNN ROIs (including PPA), and occipital regions were not specifically active during navigation in either SC or EB (except for OPA). This discrepancy might be due to the fact that, in previous studies, the navigation task was contrasted either with resting periods^39^ or scrambled tongue stimulation^38^, which require lower degrees of cognitive control. Here, using a demanding matched-control task with the exact same stimuli (numbers) but requiring a different, non-spatial computation (arithmetic), we did not find significantly greater activity in the ventral occipital cortex of EB. Crucially, when comparing navigation vs. rest, we could see that the occipital cortex was more active in EB than in SC (See Supplementary Figure S6), in line with previous reports^38, 39^. Nevertheless, such activation cannot be interpreted as strictly navigation relevant. The only brain region that was differentially and specifically activated during navigation between sighted and blind was the right inferior parietal cortex (IPC), a region of interest known to be important for egocentric spatial representations^47^. Although this effect was somewhat weak in our Nav-Math experiment, the fact that activity in this region, in EB, predicts navigation accuracy and the neural geometry in the entorhinal cortex suggests a functional role of IPC during spatial navigation.

This brings us to our second main finding: that early blindness restructured the geometry of entorhinal cognitive maps during the Clock-Navigation task. Our results show that, whereas 6-fold symmetry was detected in SC left entorhinal cortex, replicating previous imagined navigation results^51, 52^, we did not detect a similar 60° modulation of the BOLD signal in EB. On the contrary, we found the emergence of 90° sinusoidal modulation (4-fold) in EB EC, which was not detected in SC. What is the reason behind this difference across groups and the emergence of the 4-fold symmetry in EB? We can provide only a speculative explanation for this phenomenon, which is, however, in line with recent findings by other laboratories and corroborated by our own analyses. We think that 4-fold symmetry may derive from the extensive use of an egocentric perspective by blind people during navigation. Egocentric coding of the clock space would increase the reliance on the main axis of the clock, modulating the geometry of grid-cell firing fields and giving rise to a 4-fold symmetry. We will first discuss evidence supporting the link between 4-fold symmetry and the alignment with environmental axes, as well as the modulation of the entorhinal grid signal by the adoption of an egocentric perspective. Then, we will consider the details of the mechanism that could give rise to the 4-fold symmetry seen in our task.

The emergence of 4-fold symmetry in the human EC was recently reported in two fMRI studies^64, 65^ and was generally associated with an anchoring toward the cardinal axes (north-south; east-west) of the environment. He and colleagues (2019)^64^ tested participants in a virtual-reality task, in which they navigated either in an open field or in a hairpin maze. Reflecting the modulation of grid-cell firing fields when rodents navigate a similar maze^80^, a 90° rotational symmetry aligned with the main axis of the maze (north-south, east-west) emerged in humans during virtual maze navigation in fMRI^64^. We found a similar anchoring of the 4-fold entorhinal map of EB, despite the absence of barriers in our clock environment (See Fig. 3I), although in our case, grid orientations clustered with a 30° shift from the clock axes that are canonically considered as the main reference (6-12 and 3-9). In another experiment, Wagner and colleagues^65^ found the emergence of 4-fold symmetry when participants had to retrace the path of an avatar that they observed navigating in the previous trial. Similar to our clock experiment, the environment was circular and without barriers. The authors attributed 4-fold symmetry to the possibility that “cardinal directions might act as a mental coordinate system, allowing us to compare other movement directions with these major axes” (^65^, p. 11). Although the emergence of the entorhinal 4-fold symmetry could, in principle, be due to different mechanisms in these different experiments, a common factor could be the adoption of an egocentric reference frame during navigation. Indeed, the presence of barriers that partially occlude the view of the entire environment, together with the possibility of performing only 90° turns, could have fostered the adoption of egocentric navigation strategies in the hairpin maze^64^; Similarly, the task of imitating the spatial movements of a previously seen avatar^65^ may have enforced an egocentric strategy based on the relative movements of the avatar from a body-centered viewpoint. Interestingly, there is evidence that enhancing participants’ self-perception and body awareness during navigation decreases the stability of 6-fold symmetry in the entorhinal cortex and increases activity in the right IPC^66^, which, notably, is involved in egocentric spatial coding^19, 47, 61^. Moreover, in the experiment by Wagner and colleagues^65^, the right IPC activity during the observation of the moving avatar predicted the accuracy during the subsequent path-retracing. Thus, it appears that right IPC activity is associated with both distorted entorhinal geometry and navigation accuracy in tasks that encourage egocentric spatial processing^65, 66^. This state of affairs is in keeping with our analysis showing that right IPC activity correlates, in EB, both with the strength of the 4- fold symmetry in EC and with navigation accuracy in imagined and real-life navigation.

In sum, as it emerges from our surveys (Fig. 4B), congenitally blind people relied more on an egocentric coordinate system during everyday navigation^35, 37^. Indeed, constructing an allocentric map without vision – precluding the possibility of perceiving several locations or comparing different distances “at a glance” – is not easy (although certainly not impossible^20, 21^). The reconstruction of the environment layout in the blind is necessarily more diachronic and sequential than in the sighted^34^. This enhances the reliance on a first-person perspective and on route-based knowledge grounded in the simulation of successive turns (independently from the global orientation of the environment). It has been shown that egocentric coding increases the dependency on the main axes of the environment to establish item positions and moving directions, which are computed as an egocentric bearing from a given axis aligned with the “point of view” from which the environment is typically experienced^81–85^. The strategy reported by most blind participants – rotating the clock in order to have the starting/ending point in front of them – implies the calculation of the direction of movement as a function of egocentric rotation (bearing) from the main sagittal axis (6-12) from which clocks are usually experienced (Table S4). If the direction of movement is calculated by egocentric bearing from the main axes, this axis and the perpendicular one become crucial for orientation, which may re-shape grid-cell firing fields^86^, giving rise to 90° rotational symmetry.

In this paper, we have shown that the human navigation network is largely resilient to early visual deprivation. Both early-blind and sighted people selectively activate the same set of regions during imagined navigation (compared to an arithmetic task), including occipital regions such as the OPA, usually considered to be highly visual^57, 87, 88^. However, we also showed different neural geometries in the entorhinal cortex of sighted and early blind people, with the typical 6-fold symmetry emerging in SC and 4-fold symmetry in EB. Notwithstanding that the exact mechanisms remain unknown, we report evidence suggesting a relationship between the emergence of 4-fold symmetry and the engagement of the parietal cortex during navigation, together with the anchoring of the entorhinal map to the main axes of the environment. Although our results could, in part, be related to the specifics of the clock environment, we show for the first time how early blindness can modulate the neural geometry of entorhinal grid maps, possibly by encouraging an egocentric perspective during navigation. Future studies should investigate differences in neural geometry patterns between sighted and early blind individuals during the navigation of different types of environments^89^, as well as during conceptual navigation in non-spatial domains of knowledge^90–94^ across complementary egocentric and allocentric reference frames^19^.

## Methods

### Subjects

Thirty-eight participants took part in the MRI experiments. Nineteen participants were early blind (EB, 10 females; age: M = 37.37, SD = 6.13; two ambidextrous), and 19 participants constituted the sighted-control group (SC; 10 females; age: M = 36.21, SD = 6.44; one left-handed). Early blind participants were matched overall by age and sex with sighted control participants, with no significant age difference between the two groups (Paired sample t-test; t_18_ = −1.27, p = 0.21). Thirty-six participants took part in the path integration experiment (two SC participants did not come back to complete the experiment: 19 EB &17 SC, 8 females; age: M = 35.88, SD = 6.41; one left-handed). Nonetheless, no difference in age between the two groups was detected (Welch two sample t-test; t_33.15_ = - 0.70, p = 0.48). Early blind participants were blind since birth or completely lost vision before age 5, reporting, at most, faint light perception and no visual memories. All participants speak Italian fluently, and none reported a history of neurological or psychiatric disorders. The ethical committee of the University of Trento approved this study; all participants signed informed consent prior to the experiment and were compensated for their participation. None of the participants was excluded due to excessive head motion (i.e., the maximum head motion of all runs was no more than 3mm in translation or 3° in rotation).

#### Supplementary table 1 shows the demographic information of the early blind

#### Clock-Navigation Experiment Stimuli

Experimental paths were designed using AutoCAD (v2019), calculating the angle created by each line that connected each number on the clock, represented by the vertex of a dodecagon, to all the others. Repeated paths (e.g., from 9 to 12 and from 12 to 9), paths connecting two adjacent numbers (e.g., from 1 to 2), and those that traverse the center of the space were discarded, resulting in a set of 36 unique paths. Twenty-four path combinations were chosen, ensuring that all the degrees ranging from 15° to 360°, in steps of 15°, were represented, and each number from 1 to 12 was used as starting or ending location an equal number of times (four times per number). We computed the length of each path as the difference between the starting and the target numbers (e.g., from 9 to 12: 9 – 12 = 3), resulting in three different path lengths: (i) Short (distance between numbers: 3); (ii) Medium (distance between numbers: 4) and (iii) Long (distance between numbers: 5) and balanced it within each run so as to have six short paths; six long paths and 12 medium paths.

Instructions were delivered auditorily to participants (48000 Hz, 32-bit, mono), and each number was recorded in both a feminine voice and a masculine voice using an online speech synthesizer (https://ttsfree.com/text-to-speech/italian). The average intensity of all the auditory words was thresholded and equalized at 60 dB using Praat 6.1.01 (http://www.fon.hum.uva.nl/praat/).

#### Clock-Navigation Experiment Procedure

Participants performed an imagined navigation task in the fMRI scanner. The task was designed similarly to a previous imagined spatial navigation paradigm^51, 52^. Participants were asked to imagine themselves within a clock-like environment, with numbers positioned exactly as on a real clock, and to imagine walking from one number to another, according to instructions. Before the experiment began, we instructed participants to perform the shortest and straightest path possible to arrive at the end point, avoiding walking along the perimeter of the clock or stopping or turning in the center. Each trial concluded with a question about the position of a target number in the space as compared to the participants’ imagined position at that moment, as shown in figure 1A. Prior to the fMRI session, participants performed behavioral training (4 blocks, 96 trials) to familiarize themselves with the task.

The behavioral training was conducted until participants reached a performance equal to or greater than 80% for two blocks consecutively. The fMRI session consisted of 8 runs of 24 trials each (192 trials). In half of the runs, the chosen path combination (see above) was presented in the standard configuration (namely, “Forward” runs). For the other half, participants were guided in the opposite direction (“Backward” runs; e.g., Forward runs: from 8 to 2 – Reverse runs: from 2 to 8). Forward and backward runs were presented interleaved.

Participants heard two numbers (0.5s each, separated by 0.5s), recited by a female voice, that corresponded to the starting and ending points. After the ending point instruction, they were asked to begin imagining walking across the clock for a variable amount of time between 4 and 6 seconds, at the end of which, a third number (target) was recited (0.5 seconds) by a male voice. Participants were asked to decide whether the target number was situated to the left or the right in relation to the arrival position (4.5 – 6.5s). The press of a button with the right-hand middle finger indicated the number position was on the left; with the right-hand index finger to indicate the number was situated on the right. Participants were trained to answer before a sound cue played 2 seconds after the end of the target number instruction. This condition was introduced to push participants to navigate the environment during the imagination time window, reducing the possibility that they start imagining only after having heard the target number. The experimental paths were arranged so that for two consecutive trials, the previous ending point corresponded to the next trial’s starting point; when this condition was violated, participants heard a “jump” (In Italian, “salto”, 0.5s) instruction, which indicated that the starting point of the new trial would be different from the ending point of the previous one. They were given 4 seconds after the jump instruction to reorient themselves according to the new starting position. The target numbers associated with each trial were pseudo-randomized across runs, while making sure that they were different from the numbers used as starting and ending positions and were not adjacent to the ending point. Moreover, the positions of the target numbers in space were counterbalanced within each run so that they appeared an equal number of times (12) on the left and on the right side of the space. Target numbers were also counterbalanced across runs in such a way that, for the same trajectory, in half of the runs, the target number was situated on the left and the other half of the runs on the right.

During the entire duration of the experiment, participants were blindfolded and received instruction through MRI-compatible earphones using Psychtoolbox 3.0.14 (http://psychtoolbox.org/). Answers were given using an MRI-compatible response box connected to the testing PC. Stimuli presentation in the behavioral training, as well as the fMRI experiment, were displayed using Matlab releases 2014b (Behavioral Training) and 2017b (fMRI).

#### Nav-Math Experiment Stimuli

Pairs of numbers (i.e., paths for the navigation blocks) were chosen to be identical between math and navigation tasks.

Only the order of presentation of each combination was shuffled across tasks and blocks. Twenty combinations were pseudo-randomly selected out of the 36 available (see above) to make sure that each path was presented only once and not repeated backward. Each number between 1 and 12 was presented at least once and a maximum of six times to avoid number oversampling. Space resolution was not taken into account in the construction of the navigation blocks, as it was not a crucial feature for univariate analysis contrasting navigation and mathematics. Stimuli recording criteria were the same as those used for the main experiment (see above).

#### Nav-Math Experiment Procedure

During the Nav-Math experiment, participants were asked to complete a navigation task and a mathematic task. The navigation task was similar to the one in the Clock-Navigation experiment (see above). Participants heard three different numbers consecutively, referring to a starting point, an ending point, and a target number, respectively. Each number was played for 0.5 seconds, interleaved by 0.25 seconds of silence. Right after the third number was recited, participants had 3 seconds to decide whether the target number was on the left or on the right of the implied trajectory. Participants were required to answer as quickly and accurately as possible by pressing two keys using the index and middle fingers of the right hand.

At the beginning of the mathematic task, participants heard a target number (0.5s) for the entire block. Then, each trial in the block had the same structure as in the navigation task. In each trial, three numbers were played (0.5s each, separated by silence of 0.25s). Participants had to sum the first two numbers, subtract the third number from the sum, and then decide within 3 seconds whether the resulting absolute value was greater or smaller than the target number they had heard at the beginning of the block. Participants pressed buttons with their right-hand index finger if the result of the arithmetic operation was less than the target number and with their right-hand middle finger if it was greater. Left/Right and Greater/Smaller responses were counterbalanced within each block (10 times each condition). In both tasks, the first two numbers were recited in a feminine voice, whereas the third number was recited in a masculine voice. The Nav-Math experiment consisted of 4 blocks (2 navigation blocks and 2 mathematic blocks) of 20 trials each, interleaved, and counterbalanced across participants. Prior to each block, instructions were played as to which task would be performed. There were 15 seconds of silence between blocks, allowing the BOLD signal to decay.

Blindfolded participants underwent two runs of the Nav-Math experiment (80 trials for each task). Before participating in the MRI session, all participants performed a brief behavioral training (2 blocks for each condition, 40 trials per condition) to be familiarized with the experimental design.

Audio stimuli were delivered with MRI-compatible headphones using Psychtoolbox 3.0.14 (http://psychtoolbox.org/), and button presses were recorded using an MRI-compatible response box. Behavioral training and the fMRI sessions were performed using Matlab releases 2014b (Behavioral) and 2017b (fMRI).

#### MRI data collection

Echo-planar images (EPI) were acquired with a 3T Siemens Prisma scanner using a 64-channel coil. Functional images were acquired with the following parameters: Field of View (FoV) = 200mm; Voxel Size = 3×3×3mm; Number of slices: 66; Time Repetition (TR) = 1000ms; Time Echo (TE) = 28ms; Multi-band acceleration (MB) factor = 6 and a flip angle of 59°. Signal loss in the medial temporal lobe region was addressed by tilting the slice of 15° compared to the anterior-posterior commissure line (ACPC, direction: anterior edge of the slice towards the check). Moreover, to avoid slice group interference, we made sure that the ratio between the number of slices and the MB factor was equal to an odd number^95^ (66/6 = 11). Gradient-echo Field maps were acquired for distortion correction of the functional images using the following parameter: FoV = 200mm; Voxel Size = 3×3×3mm^3^; TR: 768ms; TE = 4.92 and flip angle of 60°. These images were acquired by adding a fat band to avoid the presence of wrap-around artifacts in the images. In addition, a Multi-Echo MPRAGE (MEMPRAGE) sequence was used to acquire a T1-weighted structural image for each participant with the following parameters: FoV = 256mm; Voxel Size = 1×1×1 mm; TR = 2530 ms; TE1 = 1.69ms; TE2 = 3.55ms; TE3 = 5.41ms; TE4 = 7.27ms and a flip angle of 7°.

#### fMRI data pre-processing

Images underwent standard preprocessing procedures using Statistical Parametric Mapping (SPM12 https://www.fil.ion.ucl.ac.uk/spm/software/spm12/) in Matlab 2020a. Field maps were used to calculate the voxel displacement matrix to reduce distortion artifacts. Functional images were spatially realigned and corrected with the voxel displacement matrix using the ‘Realign and Unwarp’ tool in SPM. Structural images (T1) were realigned to the mean of functional images, and the functional images were normalized to the Montreal Neurological Institute (MNI) space using unified segmentation methods. Images were smoothed with a 6 mm full-width-at-half-maximum (FWHM) spatial kernel. Whole-brain analyses were implemented on the smoothed images in the MNI space; ROI analyses in the EC were implemented on the unsmoothed images in the native space.

#### Nav-Math Experiment: Whole brain analysis

First, five conditions were modeled in the General Linear Models (GLMs) at the first level: navigation instructions, mathematic instructions, mathematic target number, navigation task, and mathematic task. The duration of the navigation and mathematic tasks was calculated from the onset of the first number until the participants’ responses (Purple boxes, Figure 1B-C). The produced boxcar functions were convolved with the hemodynamic response function (HRF). The six-rigid head motion parameters, computed during the spatial realignment in SPM, were included in the GLMs to control for head motions. Furthermore, slow drifts in the signal were removed using a high-pass filter at 1/256 Hz. Second, we performed a non-parametric permutation analysis using the SnPM toolbox for SPM (http://nisox.org/Software/SnPM13/). One-sample t-test (EB and SC independently) or two-sample t-test (SC > EB and EB > SC) were computed on each participant’s beta maps obtained by contrasting the two tasks (Navigation > Math and Math > Navigation) and by contrasting each task with the resting state periods (Navigation > Rest and Math > Rest). An Explicit mask of the main brain areas involved in navigation processing was set in the model. The HNN ROI was obtained by searching for the term ‘navigation’ in Neurosynth (https://www.neurosynth.org/) which integrated brain activation from 77 studies (cluster of activation thresholded at p_FDR_ < 0.01). We ran 10000 permutations, and no variance smoothing was performed. A voxel-level FDR corrected p = 0.05 was set as the threshold for multiple comparisons. In addition, group-level analysis using the same data reported above was performed on unmasked images, allowing exploratory whole-brain analysis. Lastly, small volume correction (SVC) analysis was performed on beta maps resulting from the group and task contrast ([Navigation > Math] × [EB> . SC]) to assess possible group difference in the parietal cortex with the voxel-level significant threshold at p < 0.001 and the cluster-level significant threshold at p_FWE_ < 0.05.

#### ROI Definition

Subject-specific entorhinal cortex in the left and right hemispheres were cytoarchitectural defined and converted from surface space to volume space using the ‘mri_convert’ function in Freesurfer (v7.1.1, https://surfer.nmr.mgh.harvard.edu/). The obtained masks were co-registered to the mean functional image to obtain the same spatial resolution (3×3×3 m^3^). On average, sighted controls’ left EC included 116 voxels and right EC 96 voxels, and early blind’s left EC included 106 voxels and right EC 93 voxels. Analyses were conducted both using individual hemisphere EC masks and bilateral EC masks combining both hemispheres (Nau et al., 2018).

#### Clock Navigation Experiment: Grid-like signal analyses

Four-way cross-validation has been applied to quadrature filter analysis previously used to detect grid-like coding in humans’ EC during spatial navigation tasks^48, 51, 96^. The eight experimental runs were divided into 4 partitions of 2 runs each, combining a ‘forward’ and a ‘backward’ run (see above). For each iteration, three partitions (6 runs) were used in GLM-1 to estimate the preferred grid orientation, and the remaining partition (2 runs) was used in GLM-2 to test the strength of the grid-like representation in the EC. The process was iterated until each partition was used for GLM-1 (Estimate) three times and for GLM-2 (Test) once (4 iterations). Within each partition, trajectories’ degrees were equally represented. In both GLMs, the signal was high-pass filtered at a threshold of 1/128 Hz to remove the slow drift, and the six-rigid head motion parameters computed during the spatial realignment in SPM were modeled as nuisance regressors. Both GLMs also included two regressors of no interest: the ‘jump instruction’ period and the period computed from the onset of the ‘target number instruction’ event until participants’ responses.

More specifically, GLM-1 was first used to estimate the preferred grid orientation for each participant in their native space. Imagined navigation periods (purple box, Figure 1A) were modulated by two parametric regressors: *cos(6θ_t_)* and *sin(6θ_t_),* where θ_t_ indicates the trajectory’s angle at trial ‘t.’ The factor ‘6’ indicates that six-fold symmetry rotational periodicity was tested. Resulting beta maps, β_1_ and β_2_, were used to calculate each voxel’s grid orientation as:

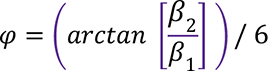

Participants’ preferred grid orientation was calculated as a weighted mean across all the voxels within the EC ROI. The weight of the mean reflected the amplitude of the voxel response calculated as follows:

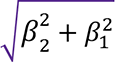

Second, the remaining partition was used in the GLM-2 to test the emergence of a 60°sinusoidal modulation of the BOLD by calculating the cosine of the trajectories’ angle performed by participants realigned with their preferred grid orientation as in:

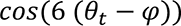

The magnitude of grid-like representation in each participant was quantified by averaging the parametric regressors estimated in the GLM-2 across all the iterations. Finally, to better visualize the effect, we divided the trajectories’ angles into 12 bins aligned with each participant’s mean grid orientation (60° modulo 0) or misaligned (60° modulo 30) and averaged the corresponding regressors across participants (Figure 4B middle). All the analyses were conducted on the periodicity of interest (6-fold) as well as on control models (4-, 5-, 7-, 8- Fold symmetries)

Finally, we modeled a group-level GLM to conduct an exploratory whole-brain analysis. One-sample t-test (independently for SC and EB) was conducted on beta maps obtained in the GLM-2 to investigate which brain areas were sensible to 6-fold symmetry.

#### Path Integration Experiment procedure

The path Integration behavioral experiment was constructed following ^96^ experimental design. Differently from the classical path integration tasks, where participants were asked either to estimate the distance or orientation of a target point from a single location, this experimental design required participants to both estimate distance and orientation of a target point from two different locations in the space. The use of multiple locations enabled the collection of a greater amount of data points, which would provide a more accurate calculation of the PI errors. Moreover, distance and orientation measurements acquired from different points in space might reduce the biases in the estimations that could arise when participants need to provide a single answer at the end of the walked path^96^. Here, we used half of a tennis field (10.97 × 11.88 m) to construct eight unique paths following the field’s predefined lines (Figure S3). In Each path, two stopping points were introduced with distances from the starting location ranging from 4m to 14m and angles from 37° to 122°.

Participants were blindfolded before the entrance into the experimental environment. They were provided with a compass attached to a string around their necks and a cardboard stick in their hands during navigation. Before the experiment started, they received verbal instructions about the task, and once they confirmed to have understood all the instructions, the experiment started. During the experiment, participants placed both hands on the stick. The experimenter stood in front of them, with a hand on the cardboard stick, and led them to walk along the path. Once reached the first stopping point, the experimenter asked participants to first estimate in meters and centimeters the Euclidean distance between their actual position and the starting point and then to point toward the direction of the starting position using the compass. The answers were transcribed by the experimenter. Participants completed eight paths, repeated twice (16 paths), and none of them reported to have recognized the repeated paths. At the start and the end of the experiment, participants completed two ‘standardization paths’ used in the analyses to correct their distance estimations (see the ‘Path Integration analyses’ paragraph). They were asked to walk two straight paths of 5m and 10m and to guess the walking distance in meters. Moreover, at the end of the experiment, they were asked whether they had heard any external cues that helped them to orient in the space during the task, but this was not the case for any of the participants.

Supplementary table 2 reports the list of the performed paths with their associated measurements.

#### Path Integration analyses

As for the experimental design, analyses were conducted following^96^. Participants’ distance estimations were corrected using the ‘standardization paths’ data to reduce the biases in the computation of the error not directly ascribable to the participants’ path integration ability but rather to their tendency to underestimate or overestimate distances in real life. First, the participants’ answers obtained at the beginning and the end of the experiment in the ‘standardization paths’ were averaged for each specific distance, 5m and 10m. Subsequently, we calculated the correction factor (C_f_) for both distances as follows:

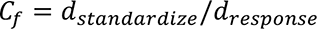

Where d_standardize_ was the actual length of the path, and d_response_ was the participants’ answer. Second, the computed correction factors were multiplied by the participants’ estimated distances during the task. Reported distances from 4m to 7.5m were corrected using the 5m C_f_, and those greater than 7.5m using the 10m C_f_. Corrected distances (d_corrected_) and orientations (Ori) estimations of each stopping point were combined to calculate the x and y coordinates of the presumed starting position:

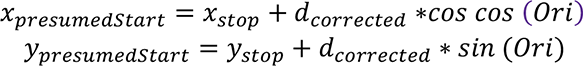

Where x_real_ and y_real_ corresponded to the real coordinates of the starting position of each trial. In order to compute the path integration error of each stopping point, the Euclidean distance between the stopping point and the presumed starting point estimated at the previous stopping point was calculated (PI Error). For the first stopping point, the previous presumed starting position corresponded to the real starting position of the path. This procedure allowed us to calculate PI independently for each path segment, reducing the possible biases produced by a cumulative computation of it. Lastly, the calculated errors were averaged across stopping points and trials, and the performance was computed as follows:

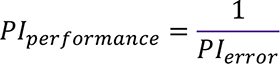

#### Questionnaires

Participants were asked to fill out two questionnaires after the fMRI experiment: one self-made questionnaire investigating which strategies they were using to navigate within the clock environment (See supplementary table 4) and a self-report questionnaire aimed to investigate participants’ navigation strategies in their everyday life^68^. The scores from the QOS were obtained by summing the points of specific questions following the guidelines provided by the authors (‘Navigation Confidence’: question 3 + question 4+ question 5s+ question 10 + question 11 + question 13; ‘Route knowledge’: q5r + q6r; ‘Survey knowledge’: q5s + q6s + q9_sp_ - 9_verb_; Pazzaglia et. al., 2000). The ‘dRS’ scores were computed by subtracting the ‘Survey knowledge’ score from the ‘Route knowledge’ score so that a positive value would indicate participants’ tendency to rely on more egocentric strategies during their everyday life navigation, and negative values would have indicated the opposite.

#### Statistical analyses

Statistical analyses were conducted using SPM and SnPM in Matlab 2020a for whole brain analyses and R (v4.2.2) for ROI analyses. The distributions of the grid orientations (Fig. 3I) were assessed using the function in the CircStat toolbox in Matlab 2020a. Results of both the whole-brain and the ROI analysis were computed using one sample t-test to investigate single-group effects and a two-sample t-test to investigate the effect between groups. The Wilcoxon-Signed-Rank test was used if the data obtained from the ROI analyses were not normally distributed. The normal distribution of the estimated parameters was checked using the Shapiro-Wilk test.

We computed Pearson’s product-moment correlations to assess the relationship between parietal cortex activity and fold symmetries or path integration performance. The correlations with brain activity in the EC were conducted using the 4-fold beta estimates in the bilateral EC of both SC and EB, whereas the correlations with 6-fold symmetry were conducted on the left EC beta estimates for SC and the bilateral EC beta estimates for the EB. The significance threshold was set at p < 0.05 two-tailed unless specified. The behavioral and the ROI results were represented using box-plot and raincloud plots, where the boxes indicate the interquartile Range (IQR), meaning the datapoints included between quartile 1 (Q1, lower quartile, 25^th^ percentile) to quartile 3 (Q3, upper quartile, 75^th^ percentile), the horizontal black line indicates the median of the values of the sample, and the whiskers indicate the distance between the upper and lower quartile to highest and the lowest value in the sample. Moreover, in raincloud plots, the distribution of the data is represented by a density curve beside each box. Asterisks upon the graphs represent significance level as follow: * p < 0.05; ** p < 0.01; *** p< 0.001

## Acknowledgments

This work was supported the by the European Research Council (ERC-StG NOAM - 804422 awarded to R Bottini). We are thankful to our blind and sighted participants for their collaboration. We are furthermore grateful to Jorge Jovicich, Nicola Pace, Stefano Tambalo, Manuela Orsini and Ilaria Mirandola for technical assistance in developing fMRI acquisition sequences. Finally, we thank all the members of the BottiniLab for the. Insightful discussion about this project.

## Authors contributions

Conceptualization, F.S., Y.X., R.B., M.S.; Methodology, F.S., and Y.X.; Investigation, F.S., and M.S.; Formal Analysis, F.S., and Y.X.; Writing – Original Draft, F.S., Y.X., and R.B.; Data Curation, F.S.; Supervision, R.B.

## Data availability

Further information and requests for data sharing should be directed to and will be fulfilled by the Lead Contact, Federica Sigismondi.

