## Supplementary Materials for "Altered grid-like coding in early blind people"

**Supplementary Table 1. Demographic information of the early blind and their matched sighted controls**

| EB CODE | AGE | GENDER | ONSET OF TOTAL BLINDNESS | ETIOLOGY | SC CODE | AGE | GENDER |
|---|---|---|---|---|---|---|---|
| EB01 | 51 | M | Birth | Optic nerve hypoplasia | SC01 | 44 | M |
| EB02 | 36 | M | Birth | Retinitis pigmentosa | SC02 | 32 | M |
| EB03 | 44 | M | Birth | Retinal burnt in the incubator | SC02 | 43 | M |
| EB04 | 33 | F | Birth | Microphthalmia | SC04 | 29 | F |
| EB05 | 36 | F | Birth | Premature retinopathy | SC05 | 33 | F |
| EB06 | 34 | F | Birth | Bilateral aplasia | SC06 | 31 | F |
| EB07 | 42 | F | Birth | Retrolenticular fibroplasia | SC07 | 45 | F |
| EB08 | 34 | M | Birth | Leber congenital amaurosis | SC08 | 31 | M |
| EB09 | 34 | F | 8 months | Congenital retinitis pigmentosa | SC09 | 36 | F |
| EB10 | 37 | M | Birth | Bilateral congenital anophthalmos | SC10 | 41 | M |
| EB11 | 35 | F | 2 years | Bilateral retinoblastoma | SC11 | 31 | F |
| EB12 | 32 | F | Birth | Premature retinopathy | SC12 | 30 | F |
| EB13 | 34 | M | Birth | Premature retinopathy | SC13 | 35 | M |
| EB14 | 38 | M | Birth | Optic nerve hypoplasia | SC14 | 34 | M |
| EB15 | 35 | F | Birth | Congenital retinitis pigmentosa | SC15 | 37 | F |
| EB16 | 37 | F | Birth | Premature retinopathy | SC16 | 34 | F |
| EB17 | 29 | M | 3 years | Dominant optic atrophy | SC17 | 30 | M |
| EB18 | 36 | F | Birth | Premature retinopathy | SC18 | 39 | F |
| EB19 | 53 | M | Birth | Bilateral congenital glaucoma | SC19 | 53 | M |

**Supplementary Table 2. Path information in the path integration task**

| START | FIRST STOP | SECOND STOP | DISTANCE: START-FIRST | DISTANCE: START-SECOND | θ START-FIRST | θ START-SECOND |
|---|---|---|---|---|---|---|
| N | D | O | 14.5 | 4.1 | 34.7° | 107.8° |
| I | C | A | 7.2 | 10.15 | 53.7° | 31.7° |
| G | O | Q | 7.6 | 11.5 | 50.5° | 33.7 |
| G | P | C | 10.4 | 7.2 | 37.9° | 115.5° |
| I | Q | O | 6.5 | 7.6 | 77.9° | 57.3° |
| B | P | H | 14.4 | 7.2 | 55.3° | 55.3° |
| O | D | L | 12.5 | 8.4 | 19.1° | 40.6° |
| N | I | O | 10.4 | 4.1 | 73.7° | 122.7° |

*$\theta$ is the inner angle of the performed segment (starting point – stopping point) defined as in:atan($y_{start,stop}/x_{stop}$)*

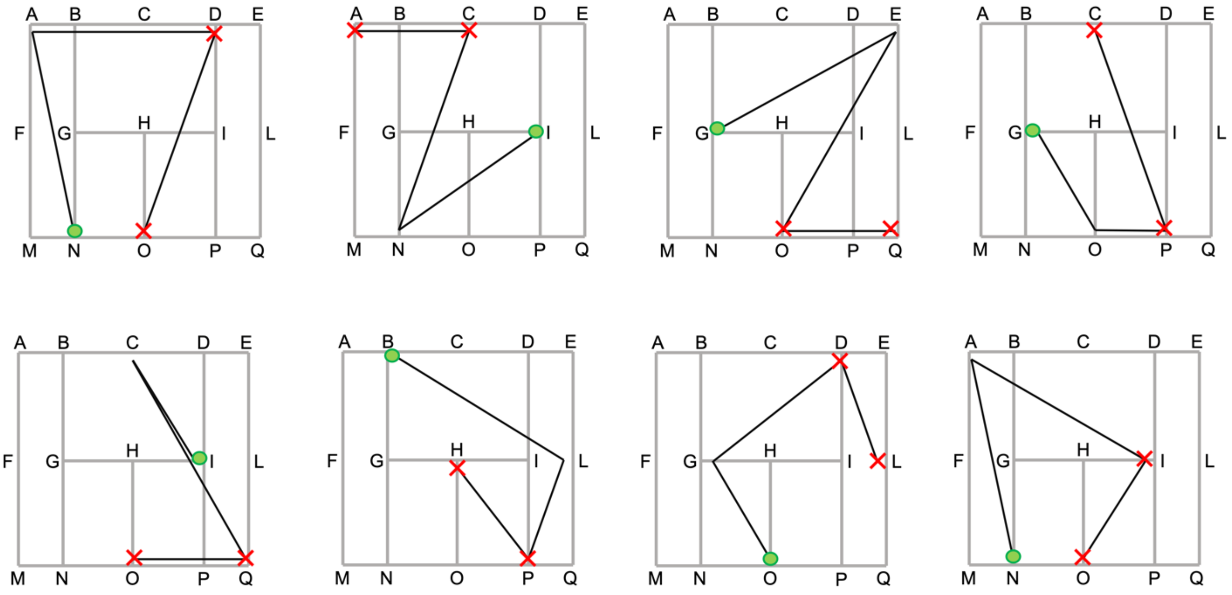


**Figure S1. Illustrations of the paths in the path integration task.**
Participants performed eight unique paths, repeated twice throughout the experiment. Each path was constituted by a starting point (green dots) and two different stopping point (red crosses). At each stopping point, blindfolded participants were required to estimate the distance and orientation of the starting point compared to their own position. The details of the paths are described in Supplementary Table 2.

**Supplementary Table 3. Brain regions more active in the navigation task than in the math task ($p_{FDR}$< 0.05)**

| Brain Regions | MNI Coordinates of the Peak Voxel | | | Peak T Value |
|---|---|---|---|---|
| | X | Y | Z | |
| **Sighted** | | | | |
| Left Middle Frontal Gyrus | -21 | 8 | 53 | 4.5 |
| Right Middle Frontal Gyrus | 24 | 2 | 53 | 5.89 |
| Left Retrosplenial Cortex | -9 | -46 | 8 | 5.29 |
| Right Retrosplenial Cortex | 15 | -55 | 11 | 5.85 |
| Left Parahippocampal Place Area | -27 | -43 | -10 | 4.14 |
| Right Parahippocampal Place Area | 27 | -37 | -13 | 4.73 |
| Left Superior Parietal Lobe/Precuneus | -12 | -70 | 53 | 7.04 |
| Left Occipital Place Area | -33 | -82 | 32 | 5.47 |
| Right Occipital Place Area | 36 | -76 | 17 | 5.54 |
| **Early Blind** | | | | |
| Left Middle Frontal Gyrus | -21 | 4 | 56 | 3.03 |
| Right Middle Frontal Gyrus | 24 | 5 | 50 | 3.70 |
| Left Retrosplenial Cortex | -12 | -55 | 11 | 3.73 |
| Right Retrosplenial Cortex | 21 | -52 | 5 | 4.62 |
| Left Parahippocampal Place Area | -33 | -37 | -16 | 3.29 |
| Right Parahippocampal Place Area | 30 | -43 | -10 | 3.09 |
| Left Superior Parietal Lobe/Precuneus | -15 | -70 | 56 | 4.89 |
| Left Occipital Place Area | -33 | -85 | 20 | 4.04 |
| Right Occipital Place Area | 42 | -79 | 11 | 4.10 |

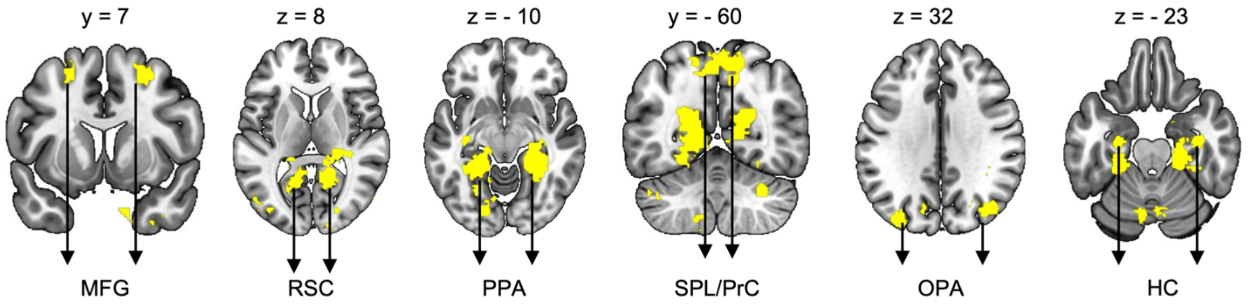


**Figure S2. The brain mask of the human navigation network**

The mask of the human navigation network consisted of (bilateral) medial frontal gyrus (MFG), retrosplenial cortex (RSC), parahippocampal place area (PPA), superior parietal lobe/precuneus (SPL/PrC), occipital place area (OPA), and hippocampus (HC). The mask was obtained from the term-based meta-analysis on *Neurosynth* across 77 studies ($p_{fdr} < 0.01$, term was "navigation"). The mask is overlapped with the MNI-152 T1 template.

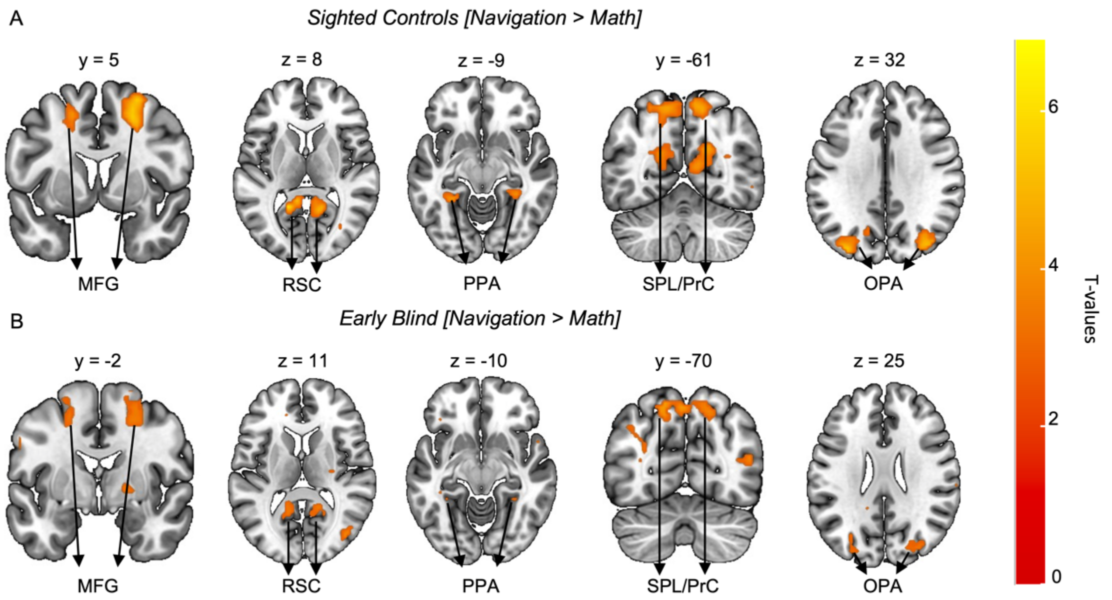


**Figure S3. Results of Navigation vs. Math contrast without the predefined mask**
Whole-brain results of the Nav-Math experiment demonstrated that both SC (A) and EB (B) activated the same network of regions (i.e., the human navigation network) during the imagined navigation of the clock space (Navigation > Math). The activations were thresholded at $p < 0.01$ uncorrected and overlapped on the MNI-152 T1 template.

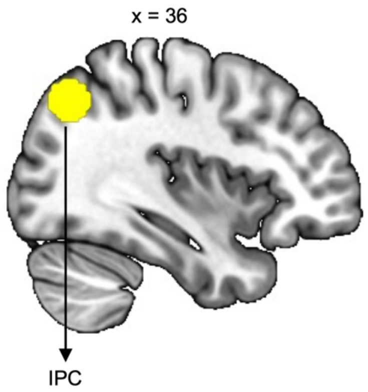


**Figure S4. EB relied more on inferior parietal cortex (IPC) during navigation than SC.**
Small Volume correction analyses were performed with a 10mm sphere around the peak coordinates from an independent study (MNI coordinates: 36/-68/44, Schindler et al., 2013), which investigated egocentric representation during an imagined navigation task. Results demonstrate a significant activation within the region of interest, suggesting that the EB compared to the SC had greater activation in the IPC during the imagined navigation task than the math task ($p_{fwe} < 0.05$; i.e., [Navigation > Math] × [EB > SC]).

**Supplementary Table 4. Strategies for imagined navigation during the Clock Navigation experiment.**

| Code | Strategies |
|---|---|
| **Sighted** | |
| SC01 | Thinking about the path I need to perform |
| SC02 | Divide the clock in two halves by tracing a line from the first to the second point |
| SC03 | Walk through the clock from the first to the second number |
| SC04 | Bird-view of the clock to locate the starting and ending position |
| SC05 | Walk through the clock form the starting to the ending point |
| SC06 | Divide the clock space with a line |
| SC07 | Visualize the clock from the starting point |
| SC08 | Walk through the clock |
| SC09 | Rotate the clock space according to my position |
| SC10 | Rotate the clock space according to the starting point position |
| SC11 | Bird-view of the clock |
| SC12 | Divide the clock in two halves from the starting to the ending point |
| SC13 | Walk through the clock |
| SC14 | Bird-view of the clock space tracing a line from the starting to the ending point |
| SC15 | Imagine the path to perform within the clock |
| SC16 | Walk through the clock |
| SC17 | Imagine walking from a number to the other |
| SC18 | Walk through the clock |
| SC19 | Imagine walking from a point to the other of the space |
| **Early Blind** | |
| EB01 | Trace a line from a point to the other |
| EB02 | Rotate the clock according to the starting point location and walked until the target point |
| EB03 | Imagine looking always at the ending point being on the starting position |
| EB04 | Imagine having the ending point in front of me |
| EB05 | Imagine two points on the clock |
| EB06 | Rotate the clock to have the starting point in front of me |
| EB07 | Imagine myself within the clock space |
| EB08 | Imagine the clock space divided in two halves accordingly to the starting and ending point |
| EB09 | Rotate the clock so to have the starting point in front of me |
| EB10 | Rotate the clock so to always face the ending point |
| EB11 | Walk from a number to the other |
| EB12 | Imagine the path to perform |
| EB13 | Rotate the clock to have the starting point in front of me |
| EB14 | Rotate the clock to have the starting point in front of me |
| EB15 | Rotate the clock to have the starting point in front of me |
| EB16 | Imagine the ending point in front of me |
| EB17 | Divide the clock with a line |
| EB18 | Imagine the ending point in front of me |
| EB19 | Imagine the ending point in front of me |

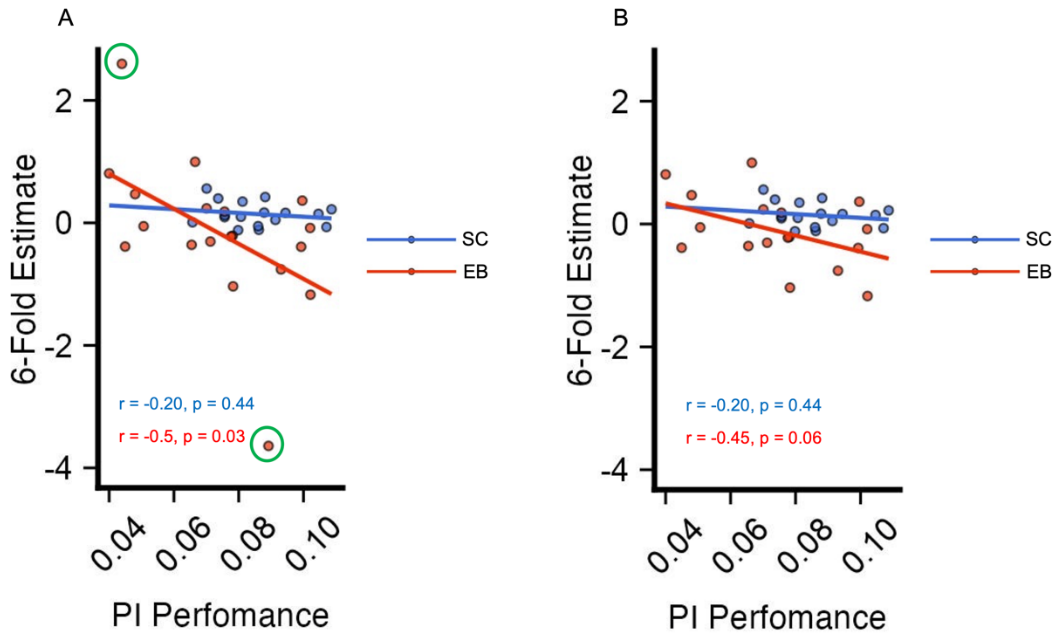


**Figure S5. Path Integration (PI) ability negatively correlated with 6-Fold symmetry in Early Blind participants.**

(A) 6-Fold symmetry estimates negatively correlated with PI Performance in early blind participants, suggesting that sighted-like representation of space might be dysfunctional for navigating without vision (Pearson's product-moment correlations; r = -0.5, p = 0.03). However, this correlation might be biased by the presence of two outliers (green circles) in EB population. No significant correlation was found in the sighted controls group (r = -0.19, p = 0.44). (B) Removing the outliers in the early blind group weakened the negative correlation between 6-Fold symmetry estimates and PI performance (r = -0.45, p = 0.06).

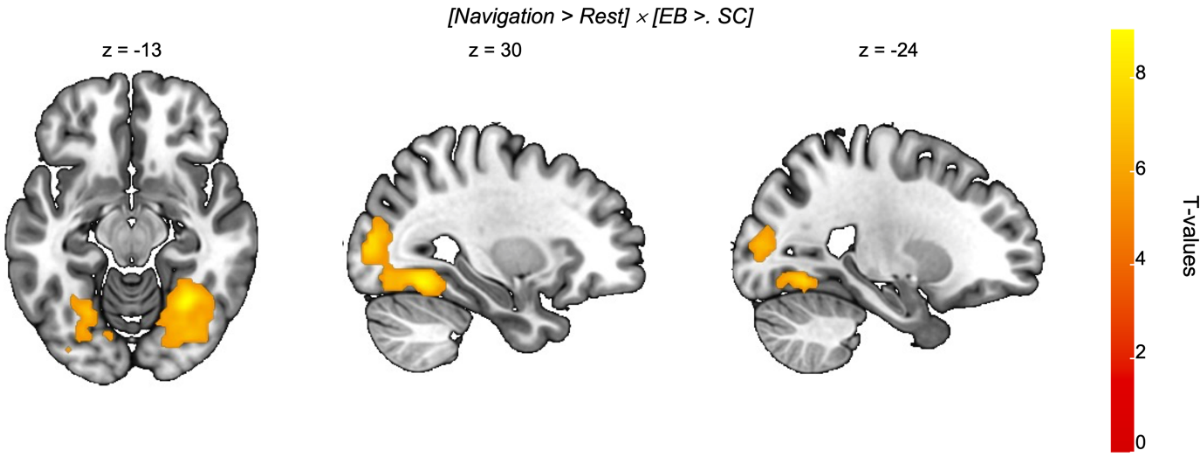


**Figure S6. Occipital cortex was more activated in EB than SC.**
Although no difference between SC and EB were detected when the navigation task was compared to the math task, when comparing the navigation task against rest ([Navigation > Rest] × [EB > SC]), we could find the emergence of clusters of activity in several occipital areas among which bilateral inferior occipital gyrus; occipital fusiform gyrus; fusiform gyrus and lingual gyrus in EB more than SC, which is in line with previous results (Gagnon et al., 2012; Kupers et al., 2010. The activations were thresholded at $p_{FWE} < 0.05$ and overlapped on the MNI-152 T1 template).
